# Changes in housing and diet combined increase fecal Oxyurid load in captive tokay geckos

**DOI:** 10.1101/2025.07.27.667008

**Authors:** Diana S. Gliga, Birgit Szabo

## Abstract

Reptiles are increasingly popular as exotic pets and suffer high mortality especially in the first year in captivity, yet research into their welfare remains limited. Reptiles are often infected by parasites. One common taxon is Oxyuroidea superfamily, which has a direct life cycle that promotes easy transmission in enclosed environments. Due to limited ecological knowledge, inappropriate husbandry practices are common in reptiles causing stress and increased parasite loads leading to severe health issues. Therefore, it is crucial to understand how routine captive procedures may influence parasite infections. In this study, we exposed captive bred tokay geckos (*Gekko gecko)* to two stressors common in the reptile trade, cohabitation with a novel conspecific and a change in diet. We tested their effect on fecal oxyurid output compared to a control group as well as their combined effect. We found that a single stressor had no effect on fecal parasite load while both combined did significantly increase parasite load. However, we did not find a detectable change in the lizards’ general condition. Our study shows that seemingly minor changes in housing and husbandry can exert stress and increase parasite load in tokay geckos. Further studies are needed to determine which other procedures (e.g. confinement, transport, novel environments) affect health and when combined could lead to more severe changes in health in reptiles.

**Summary:** Many reptiles are infected with parasites that can lead to health issues. Routine husbandry procedures such as new conspecifics and diet are stressful and lead to a higher parasite load in geckos.

## Introduction

Reptiles are becoming more and more popular as exotic pets around the globe. In 2022, it was estimated that 1.5 million individuals were kept in captivity in the UK alone, an increase of 60% from 0.9 million estimated in 2016 (PFMA, 2016; 2022). Despite this large number of individuals being held in captivity, research into reptile welfare is still lacking behind compared to other taxonomic groups such as mammals (Doody, 2023). Especially during the first year in captivity, reptiles suffer a heightened risk of mortality (Robinson et al., 2015). This increased mortality risk is caused by insufficient knowledge regarding ecology and natural history of different species leading to suboptimal captive conditions and husbandry (Warwick, 2014; Wilkinson, 2015). Furthermore, many captive reptiles harbor several internal parasites (Arabkhazaeli et al., 2018; Hallinger et al., 2019; Rataj et al., 2011) which, under inadequate management, could contribute to the increased mortality risk. Transport, handling, new environments and unfamiliar conspecifics are all common stressors (Langkilde and Shine, 2006) within the pet reptile trade that could negatively affect immune function and lead to a parasite infection getting out of hand (e.g. Hallinger et al., 2018). Therefore, a better understanding of which routine procedures cause stress and exacerbate parasite infections is needed.

During the co-evolution with its host a parasite can reduce its virulence, as is the case of the Oxyuroidea superfamily which have the broadest host distribution of the nematodes and are highly host species-specific (Adamson 1994). They are also among the most prevalent nematodes found in captive reptiles (Hallinger et al., 2018) including geckos (Amaral et al., 2021) and bearded dragons (Schmidt-Ukaj et al., 2017). Reptile oxyurids belong mainly to the family Phayngodonidae (Jacobson and Garner 2020). These produce two types of eggs: thin-shelled eggs that hatch in the intestine and are readily auto-infective (endogenous life cycle) and thick-shelled eggs that require excretion in the environment to become infective (exogenous life cycle) (Adamson 1994). The direct life cycle and the high resistance of the environmental stages (eggs) favour the transmission in captive settings. Furthermore, bedding and enrichment material can be difficult to keep clean or treat increasing the chances of reinfection. Oxyurid infections are usually well tolerated in healthy reptiles (Brosda, 2013) and the species-specific parasite-host relationship is also believed to have indirect beneficial effects such as improved absorption of nutrients during digestion, particularly in herbivores (Brosda, 2013). However, high burdens might lead to intestinal obstruction (Schmidt-Ukaj et al., 2017) and in extreme cases end deadly, as demonstrated in pet tortoises (Hallinger et al., 2018). Consequently, especially in the period after the acquisition of a new pet reptile, stressors need to be avoided. However, which specific procedures lead to changes in parasite load is unclear.

In this study, we investigated the effect of experimental stressors such as change in housing (from single to pair housing with an unfamiliar, opposite-sex individual) and change in diet (from crickets to cockroaches) and their combined effect on the fecal parasite output in tokay geckos (*Gekko gecko*). Both scenarios occur commonly after acquisition of a pet reptile or trade by a new owner. Tokay geckos are a very popular pet species around the globe (Grossmann, 2006). They are medium sized, nocturnal, arboreal geckos that feed mostly on insects but opportunistically also feed on small vertebrate prey (Bucol and Alcala, 2013; Grossmann, 2006). They form pairs, perform biparental care and form family groups with their offspring (Grossmann, 2006). We hypothesised that the stress related to a change in housing (new environment, unfamiliar conspecific; Langkilde and Shine, 2006) or change in diet would be sufficient to cause a change in the parasite load and predicted an increase in parasite output in feces. However, each alone might not cause enough stress, and therefore, we also hypothesised that both stressors would have an additive effect and predicted a higher parasite output in individuals that experienced both a change in housing and change in diet.

## Methods

### Study animals and husbandry

We collected data from 39 adult tokay geckos (*Gekko gecko*) which either originated from different breeders across Europe (N = 11 females and 10 males) or were bred from these individuals in captivity (N = 12 females and 6 males). We used 23 females and 16 males (presence = male or absence = female of femoral pores; Grossmann, 2007) which were between 2-9 years old.

Geckos were housed in enclosures made of rigid foam slabs (female single housing: 45L × 45B × 70H cm, males single housing: 90L × 45B × 100H cm, pair housing: 90L × 45B × 100H cm; only suitable for scientific purposes) with glass sliding doors at the front and a top made out of mesh for improved ventilation. Enclosures included a compressed cork back wall with shelters hung on the back wall (hollow cork branches cut in half), cork branches for climbing and life plants for enrichment. The ground was made of organic rainforest soil (Dragon BIO-Ground) or seedling soil (various brands) on top and expanded clay as the bottom for drainage separated by mosquito mesh. Enclosures were set up bioactive and included isopods and earth worms for which we spread autoclaved red oak leaves and sphagnum moss on the soil to hide in. To provide opportunity for further thermoregulation, a heat mat (Tropic Shop) was fixed to the left outside wall of each enclosure locally increasing the temperature by 4-5°C. Furthermore, all enclosures were equipped with a UVB light (Exo Terra Reptile UVB 100, 25 W) which provided UVB during the light phase. Tokay geckos are nocturnal. To be able to work with them during their natural activity period, we kept them under a reversed 12h:12h photo period (light: 6 pm to 6 am, dark: 6 am to 6 pm; including simulated sunrise and sunset). Visibility was further improved by a red light (PHILIPS TL-D 36 W/15 RED) invisible to the geckos (Loew, 1994) during the dark phase. The change in light phase was accompanied by a gradual change in temperature from 31°C during the day and 25°C during the night simulating natural conditions. Room humidity was set to 50% but is increased to 100% for a short period of time by means of reverse osmosis water provided by an automatic rain system twice a day (30s every 12 h at 5 pm and 4 am each day). The lizards are kept across two rooms, on shelves (large enclosures on the bottom and small enclosures on the top).

Geckos were fed on Mondays, Wednesdays and Fridays with 25 cm long forceps. Before the start of data collection all geckos were fed with 3-5 adult house crickets (*Acheta domesticus*). Insects were fed with cricket mix (reptile planet LDT, which provides Vitamin D and calcium), dry cat food (various brands) and fresh carrots to provide optimal diet. A water bowl (cleaned at least once a week or more often when dirty) in each enclosure provided water ad libitum.

### Experimental set-up and procedure

At the start of the experiment, all 39 individuals were housed singly and fed with crickets. They had been in single housing for more than a year and had been feeding on crickets since arrival or birth (except when tested for food neophobia: October 2021 and April/ May 2022; Szabo and Ringler, 2023a; b). We collected a fresh (within 8h of defecation) fecal sample from all individuals during single housing between the 1^st^ and 21^st^ of December 2023 in the mornings (8:00 am to 12:00 pm). Geckos produce a single fecal “scat” every 2-4 days (personal observation). They defecate in a latrine (also called scat pile; Grossmann, 2007). Often, both individuals defecate in the same latrine (unpublished data), although not necessarily on the same day (personal observation).

In January 2024, we formed 15 stable pairs moving females into the larger enclosures of males. The remaining 9 individuals (8 females and 1 male) were kept singly for the rest of the experiment. One week after pair formation, we changed the diet of eight randomly selected pairs to cockroaches (*Nauphoeta cinerea*). Simultaneously, we switched five randomly selected individuals that were still in single housing to a diet of cockroaches. The remaining individuals were kept on a diet of crickets. We waited a week after pair formation to ensure pairs were stable and did not show any aggression towards each other. Another two weeks later, we collected a second fecal sample of all individuals between the 10^th^ and 30^th^ of January 2024. In some cases, we were able to sample feces separately from both individuals in a pair (sharing an enclosure) but for some we were unable to determine which sample came from which individual. These were pooled into one sample (two pieces of feces) per pair. Before sample collection, all tubes were treated with UV for 15 minutes. All samples were collected by the same researcher (BS) who wore new gloves to collect each sample by hand (easier to grab). All samples were frozen at -20°C 5-10 minutes after collection.

### Preliminary parasitological examination

A preliminary screening was done on three pools of fresh fecal samples to check for the presence of parasites: (1) Fecal smears were examined natively on a glass slide with a drop of saline, (2) Ziel-Neelsen staining, and (3) the combined sedimentation-flotation method (Deplazes et al., 2016). Native smears revealed few trophozoites of flagellates. No acid-fast parasite stages were seen (e.g., *Cryptosporidium* sp). The flotation slides were positive for oxyurids. DNA was extracted directly from feces using the Quick-DNA™ Fecal/Soil Microbe MiniPrep Kit (Zymo Research, USA), according to manufacturer instructions. To characterize the flagellates, a PCR that amplified the 5.8S rRNA region flanked by ITS1 and ITS2 was performed using the published ITSF and ITSR primers (Smejkalová et al., 2012). The amplicons were sent for Sanger sequencing to a commercial provider (Microsynth AG, Balgach, Switzerland). The obtained sequence matched *Simplicimonas simlis* in one direction (consensus length 145 bp, 99.3 % pairwise identity with KJ101561).

### Quantitative parasitological examination

We expressed the burden of parasites as oxyurid eggs per gram (EPG). Oxyurid eggs in reptile belong to the Pharyngodonidae, where it is believed that intra-intestinal auto-infection can occur (Adamson 1994), as opposed to the transmission strategy in oxyurids of mammals, where egg deposition occurs in the peri-anal area and infection occurs by fecal-oral route. Each fecal pellet was weighted and placed in a 10 ml weighted glass tube. The tube was filled with water up halfway and vortexed. The homogenate was drained through a double gauze on funnel in another glass tube. The tube was centrifuged at 600 g for 3 minutes. The supernatant was discarded, and the flotation solution (44% zinc chloride, density 1.320) was added with a pipetting bottle until halfway. The sediment was loosened and mixed with single use plastic Pasteur pipettes. More flotation liquid was added until the small cupola was formed. A 18 mm × 18 mm coverslip was placed on the top of the formed cupola. The tube was centrifuged at 600 g for 5 min. Afterwards the coverslip was placed on a microscope slide and the oxyurid eggs were counted. We scored the oxyurid eggs into four levels based on the number of eggs observed on the coverslip (parasite score; 0 = no oxyurid eggs, 1 = 1-200 eggs, 2 = 201-300 eggs, 3 = >300 eggs).

### Statistical analyses

All statistical analyses were run in R version 4.4.2 (R Core Team, 2025). We used cumulative link mixed models (CLM, Christensen, 2023) to analyse the effect of housing (single and pair housing) as well as diet (cricket and cockroach) and their interaction on the parasite score. Because we were unable to identify individual fecal pellets in 10 out of 15 pairs during pair housing, we calculated pair fecal mass (summing up the mass of the pellet of the male and the female) and pair parasite score (rounded up average of the individual scores) for the 5 pairs with individual measurements (individual data are included in the raw data file). In addition, we included sex as well as the fecal mass (to account for differences in pellet size) as fixed effects. To account for repeated measures and that individuals in a pair were given the same measurement during pair housing, we used animal identity and pair identity as random effects. To investigate the result of the interaction, we used Least-squares means comparisons (LSM, Lenth, 2025). We did not investigate changes in weight due to parasite infection because weight loss is slow in reptiles (personal observation, also see Moore et al., 2007), and from experience, many factors (e.g. egg laying in females, better/ worse feeding rate under different housing conditions, continuing growth of younger individuals) can have a much larger influence on weight fluctuations than the parasite infection.

### Ethical note

Our methodology followed the Guidelines for the ethical use of animals in applied animal behaviour research by the International Society for Applied Ethology (Sherwin et al., 2003). Experiments were approved by the Swiss Federal Food Safety and Veterinary Office (National No. 33232, Cantonal No. BE144/2020, BE9/2024). Captive conditions were approved by the Swiss Federal Food Safety and Veterinary Office (Laboratory animal husbandry license: No. BE4/11). During pair formation, we monitored adults closely and immediately separated them if any aggression occurred within the first 24 hours of pairing.

## Results

The overall median parasite score was 1 (IQR: 1-2). We found no evidence that parasite score differed between males and females (CLM, estimate_male_ = 0.701, std. error = 0.578, *z*-value = 1.212, *p*-value = 0.226) or that it was associated with the mass of the fecal pellet (CLM, estimate = 0.598, std. error = 0.972, *z*-value = 0.615, *p*-value = 0.539). Post Hoc comparisons showed evidence that changing both the housing condition as well as diet increased the parasite score (LSM, estimate_cockroach-cricket_ = 2.85, std. error = 1.13, *z*-value = 2.516, *p*-value = 0.012; Figure 1). However, we found no evidence that changing housing conditions from single to pair housing without changing the diet (continuing to feed crickets) influenced parasite score (LSM, estimate_pair-single_ = -0.583, std. error = 1.04, *z*-value = -0.561, *p*-value = 0.575). Similarly, we found no evidence that changing the diet from crickets to cockroaches without changing the housing conditions influenced the parasite score (LSM, estimate_cockroach-cricket_ = 1.04, std. error = 1.03, *z*-value = 1.003, *p*-value = 0.316).

**Figure 1.**
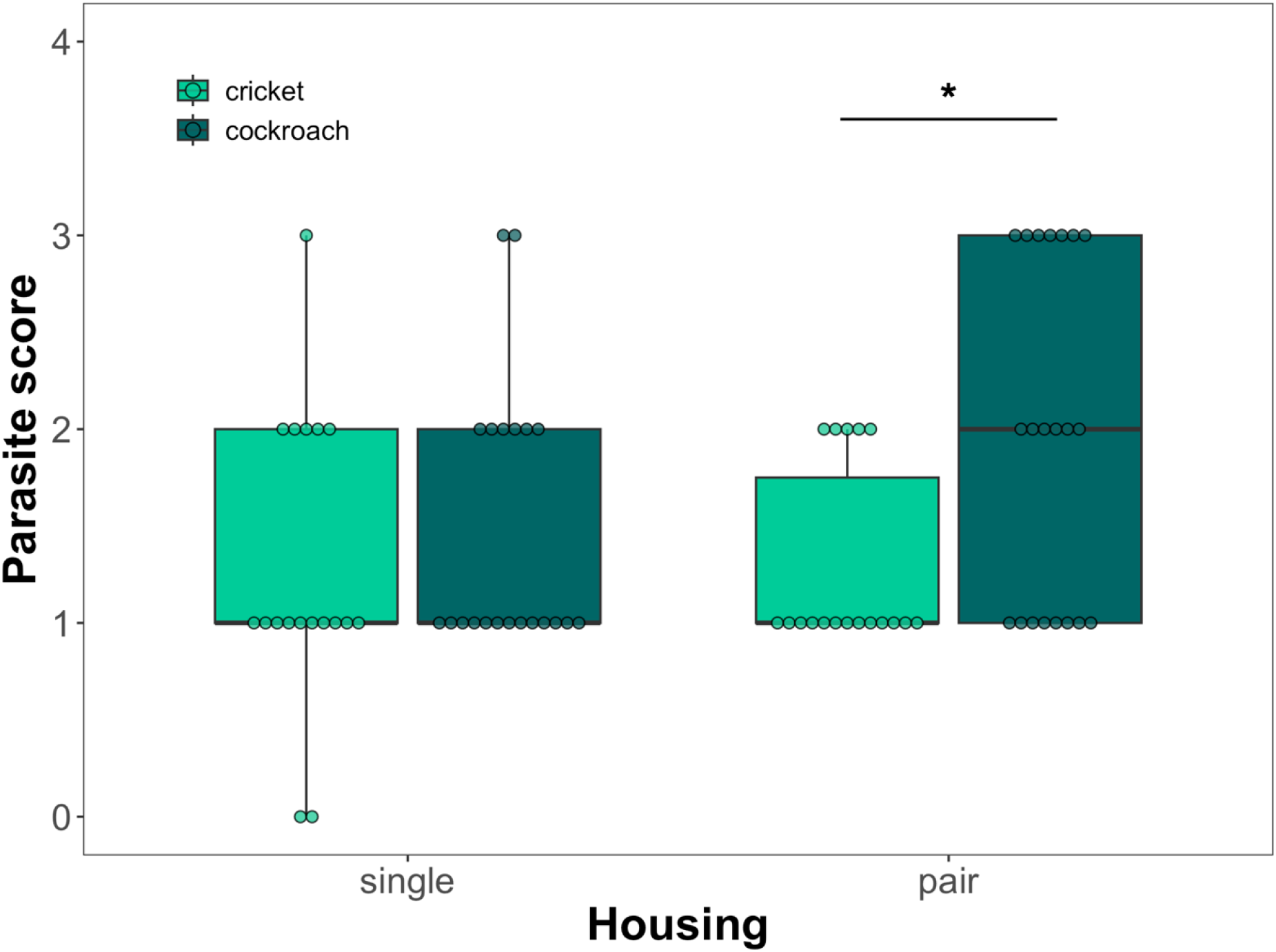
Boxplots of parasite score (0 = not detectable, 1 = 1-200 eggs, 2 = 201-300 eggs, 3 = more than 300 eggs) separated into housing treatments (single and pair housing) as well as diet treatments (cricket – light green and cockroach – dark green). Points represent individual scores. The bold line within boxes is the median, the upper box edges are the upper quartile, the lower box edges the lower quartile, the top whisker ends are the maximum and the bottom whisker ends the minimum. ^*^p < 0.05.

## Discussion

The aim of this study was to investigate if a change in diet and housing would impact the parasite output of captive bred tokay geckos. Contrary to our expectation, we found that a diet change without a change in housing and a housing change without a change in diet did not significantly impact the oxyurid output in the gecko feces. However, we found that a combination of both significantly increased the oxyurid output.

In the pet trade, reptiles often experience a number of potential stressful events including capture, transport, a new housing environment and new diet. In isolation, these events might not necessarily have a strong impact on an individuals’ health, especially when the event is short, or animals can appropriately deal with the stressor by hiding or avoidance. However, chronic stress and/ or multiple stressors together can have a more severe impact and decrease health. We find that both a change in housing and diet together increase oxyurid load suggesting that these changes are stressors that may impact lizard health. Despite this, we did not observe any noticeable change in the lizards’ body condition (weight) suggesting that the oxyurid load we measured was not severe enough to have a more detrimental effect on lizards’ health. In juvenile bearded dragons (*Pogona vitticeps)* a median oxyurid EPG as high as 14,409.25 did not lead to any loss in body condition over time (Pike et al., 2023). A further endoparasite survey in pet bearded dragon found high number of oyurids eggs (>1200 EPG) both in healthy and clinical cases (Guardone et al., 2024). Nonetheless, welfare might have still been impacted if the increased oxyurid load led to other discomforts that might have shown in a change in behaviour which we did not measure. Future studies could investigate how different oxyurid loads (before and after treatment or reinfection) impact lizard behaviour to reveal more subtle effects on welfare.

In our study, we ensured that sample collection was done systematically and as standardised as possible. For example, we made sure that the fecal sample was collected within 8 hours of defecation and only collected samples that were still fresh. Furthermore, we collected all samples in the mornings and ensured no cross contamination occurred. A previous study in bearded dragons showed that captive juveniles have 2.5x higher egg output in the afternoon than in the morning (Pike et al., 2023). If this is also true in tokay geckos needs to be investigated in the future.

Tokay gecko males are territorial in the wild and call to attract females to their territory (Grossmann, 2007). We tried to mimic this behaviour in captivity by introducing females to male enclosures. This meant that for males the change in housing included the addition of an unfamiliar female, while for females it included both an enclosure change and cohabitation with an unfamiliar male. Consequently, females might have experienced higher stress, but we were not able to reliably collect fecal samples from males and females within the same enclosure. Therefore, it is possible that females suffered a higher increase in oxyurid load compared to males, which should be investigated in the future.

In the presented husbandry our geckos appeared healthy, despite carrying oxyurids at varying intensities. The presence of the detected parasites did not affect the health status of the animals even after being exposed to stressors. Therefore, we can recommend the described husbandry system for reptile breeders and pet owners. Based on molecular data we confirmed the presence of *Simplicimonas simlis* in *Gecko gecko*. This has only been reported in one other gecko species: *Uroplatus lineatus* (Čepička et al., 2010) and a ruminant (carabao, *Bubalus bubalis kerabau*)(Dimasuay et al., 2013). Its clinical relevance is not known. Further study should explore the reptile intestinal parasitome and the delicate relationship between host and facultative parasites.

## Acknowledgements

We would like to thank Caroline Frey for her helpful comments on an earlier version the manuscript. We are also grateful to the diagnostic laboratory team at the Institute of Parasitology, Bern, for their generous provision of facilities, as well as their scientific and technical support.

## Data and resource availability

All data generated during this study and the code for analysis are available on the Open Science Framework (OSF, link for review purposes: https://osf.io/xjyfu/?view_only=46709e4b14354053aea78015b50fb2a3)

## References

Adamson, M. (1994). Evolutionary patterns in life histories of Oxyurida. Int. J. Parasitol. 24, 1167–1177. doi:10.1016/0020-7519(94)90189-9.

Amaral, C. B., Alves, A. C. C., Peroba, S. C. and Martins, I. V. F. (2021). Coproparasitologic survey of gastrointestinal parasites in a captive leopard geckos collection (Eublepharis macularius). Vet. Parasitol. Reg. Stud. Reports 26, 100617. doi:10.1016/j.vprsr.2021.100617.

Arabkhazaeli, F., Rostami, A., Gilvari, A., Nabian, S. and Madani, S. A. (2018). Frequently observed parasites in pet reptiles’ feces in Tehran. Iran. J. Vet. Med. 12, 19–25.

Brosda, A. (2013). Untersuchungen zur Infektion mit Oxyuren bei mediterranen Landschildkröten in menschlicher Obhut und ihr Einfluss auf die Entwicklung juveniler Testudo graeca. PhD thesis, Freie Universität Berlin, Berlin, Germany.

Bucol, A. and Alcala, A. (2013). Tokay gecko, Gekko gecko (Sauria: Gekkonidae) predation on juvenile house rats. Herpetol. Notes 6, 307–308.

Čepička, I., Hampl, V. and Kulda, J. (2010). Critical taxonomic revision of parabasalids with description of one new genus and three new species. Protist 161, 400–433. doi:10.1016/j.protis.2009.11.005.

Deplazes, P., Eckert, J., Mathis, A., von Samson-Himmelstjerna, G. and Zahner, H. (2016). Parasitology in Veterinary Medicine. Wageningen, Netherlands: Wageningen Academic Publishers.

Dimasuay, K. G. B., Lavilla, O. J. Y. and Rivera, W. L. (2013). New hosts of Simplicimonas similis and Trichomitus batrachorum identified by 18S ribosomal RNA gene sequences. J. Parasitol. Res. 2013, 1–5. doi:10.1155/2013/831947.

Doody, J. S. (2023). Social behaviour as a challenge for welfare. In Health and Welfare of Captive Reptiles (ed. R. J. Warwick and C. Arena), pp. 189–209. Cham: Springer International Publishing.

Grossmann, W. (2006). Der Tokeh, Gekko gecko. Münster, Germany: Natur und Tier Verlag.

Guardone, L., Marigliano, A., Mancianti, F. and Perrucci, S. (2024). Endoparasite infections in captive inland bearded dragons (Pogona vitticeps) in Italy. Pathogens 13, 443. doi:10.3390/pathogens13060443.

Hallinger, M. J., Taubert, A., Hermosilla, C. and Mutschmann, F. (2018). Occurrence of health-compromising protozoan and helminth infections in tortoises kept as pet animals in Germany. Parasites Vectors 11, 352. doi:10.1186/s13071-018-2936-z.

Hallinger, M. J., Taubert, A., Hermosilla, C. and Mutschmann, F. (2019). Captive agamid lizards in Germany: Prevalence, pathogenicity and therapy of gastrointestinal protozoan and helminth infections. Comp. Immunol. Microbiol. Infect. Dis. 63, 74–80.

Jacobson, E. R. and Garner, M. M. (eds) (2020). Infectious Diseases and Pathology of Reptiles: Color Atlas and Text, 2nd edn. Boca Raton, FL: CRC Press.

Langkilde, T. and Shine, R. (2006). How much stress do researchers inflict on their study animals? A case study using a scincid lizard, Eulamprus heatwolei. J. Exp. Biol. 209, 1035–1043.

Lenth R. (2023). emmeans: Estimated Marginal Means, aka Least-Squares Means. R package version 1.8.6-090001, <https://github.com/rvlenth/emmeans>.

Loew, E. R. (1994). A third, ultraviolet-sensitive, visual pigment in the Tokay gecko (Gekko gecko). Vision Res. 34, 1427–1431.

Moore, J. A., Hoare, J. M., Daugherty, C. H. and Nelson, N. J. (2007). Waiting reveals waning weight: Monitoring over 54 years shows a decline in body condition of a long-lived reptile (tuatara, Sphenodon punctatus). Biol. Conserv. 135, 181–188.

PFMA (2016). Pet Population Report. London, UK. 10.1086/656217.

PFMA (2022). Annual report (online) https://ukpetfood-reports.co.uk/annual-report-2022/.

Pike, C., Hsieh, S. and Baling, M. (2023). Monitoring infection load of oxyurid (Nematoda) and Isospora (Coccidia) in captive inland bearded dragons (Pogona vitticeps). Pathways Anim. Health Welf. 2, 77–90. doi:10.34074/piahw.002106.

Rataj, A. V., Lindtner-Knific, R., Vlahović, K., Mavri, U. and Dovč, A. (2011). Parasites in pet reptiles. Acta Vet. Scand. 53, 1–21.

Robinson, J. E., St. John, F. A. V., Griffiths, R. A. and Roberts, D. L. (2015). Captive reptile mortality rates in the home and implications for the wildlife trade. PLoS ONE 10, e0141460. doi:10.1371/journal.pone.0157519.

Schmidt-Ukaj, S., Hochleithner, M., Richter, B., Hochleithner, C., Brandstetter, D. and Knotek, Z. (2017). A survey of diseases in captive bearded dragons: A retrospective study of 529 patients. Vet. Med. 62, 508–515. doi:10.17221/162/2016-VETMED.

Sherwin, C. M., et al. (2003). Guidelines for the ethical use of animals in applied animal behaviour research. Appl. Anim. Behav. Sci. 81, 291–305.

Smejkalová, P., Petrželková, K. J., Pomajbíková, K., Modrý, D. and Čepička, I. (2012). Extensive diversity of intestinal trichomonads of non-human primates. Parasitology 139, 92–102. doi:10.1017/S0031182011001624.

Szabo, B. and Ringler, E. (2023a). Fear of the new? Geckos hesitate to attack novel prey, feed near objects and enter a novel space. Anim. Cogn. 26, 537–549.

Szabo, B. and Ringler, E. (2023b). The influence of partner presence and association strength on neophobia in tokay geckos. bioRxiv. doi:10.1101/2023.01.09.523205.

Warwick, C. (2014). The morality of the reptile “pet” trade. J. Anim. Ethics 4, 74–94.

Wilkinson, S. L. (2015). Reptile wellness management. Vet. Clin. Exot. Anim. Pract. 18, 281–304.

